# Neuron-derived extracellular vesicles extracted from plasma show altered size and miRNA cargo as a function of antidepressant drug response

**DOI:** 10.1101/2021.02.03.429557

**Authors:** Saumeh Saeedi, Corina Nagy, Jean-Francois Théroux, Marina Wakid, Laura M. Fiori, Pascal Ibrahim, Jennie Yang, Susan Rotzinger, Jane A. Foster, Naguib Mechawar, Sidney H. Kennedy, Gustavo Turecki

## Abstract

Previous work has demonstrated that microRNAs (miRNAs) change as a function of antidepressant treatment (ADT) response. However, it is unclear how representative these peripherally detected miRNA changes are to those occurring in the brain. This study aimed to use peripherally extracted neuron-derived extracellular vesicles (NDEVs) to circumvent these limitations and investigate neuronal miRNA changes associated with antidepressant response. Samples were collected at two time points (baseline and after 8 weeks of follow-up) from depressed patients who responded (N=20) and did not respond (N=20) to escitalopram treatment, as well as controls (N=20). Total extracellular vesicles (EVs) were extracted from plasma, and then further enriched for NDEVs by immunoprecipitation with L1CAM. EV size was measured using tunable resistive pulse sensing, and NDEV miRNA cargo was extracted and sequenced. Subsequently, studies in cell lines and postmortem tissue were conducted. Characterization of NDEVs revealed they were smaller than other EVs isolated from plasma (p<0.0001), had brain-specific neuronal markers, and contained miRNAs enriched for brain functions (p<0.0001) Furthermore, NDEVs from depressed patients were smaller than controls (p<0.05), and NDEV size increased with ADT response (p<0.01). Finally, changes in NDEV cargo, specifically changes in miR-21-5p, miR-30d-5p and miR-486-5p together (p<0.01), were associated with ADT response. Targets of these three miRNAs were altered in brain tissue from depressed individuals (p<0.05). Together, this study indicates that changes in peripherally isolated NDEVs can act as both a clinically accessible and informative biomarker of ADT response specifically through size and cargo.

## Introduction

Major depressive disorder (MDD) is a leading cause of global disability, affecting over 260 million people worldwide.^1^ Antidepressant treatment (ADT) is a first-line therapeutic intervention for MDD, however approximately 60% of patients do not respond to the first ADT prescribed.^2^ A better understanding of the mechanisms of ADT response is important for developing new avenues for treatment and to personalize pharmacological interventions.

MicroRNAs (miRNAs) are small, non-coding, single-stranded RNA transcripts that regulate the expression of messenger RNA (mRNAs) through RNA degradation or translational repression.^3^ Although several studies have implicated miRNA changes in psychopathological processes, including those associated with MDD and antidepressant drug response ^4^, most of these studies have used peripheral samples.^3^ While peripheral biomarkers are of interest in clinical settings, their capacity to reflect brain processes is debatable. The possibility of studying miRNAs encapsulated in extracellular vesicles derived from brain cells provides an exciting new avenue to study biomarkers of mental disorders and mechanisms of treatment response.

Extracellular vesicles (EVs) are released by most, if not all cell types into the extracellular space.^5^ They contain biological material from their cell of origin and can impact processes in recipient cells.^5,6^ Although different classes of vesicles have been described, exosomes (30-200nm) and similarly-sized microvesicles are of interest in psychiatry, as they can cross the blood-brain barrier and be accessed in biofluids.^6^ These important properties render them particularly good biomarkers for central nervous system (CNS) disorders. In the brain, exosomes and other EVs are important regulators of cell communication, are involved in synaptic plasticity, neurogenesis, and neuron protection among other processes.^7^ Exosomal cargo is particularly enriched in small non-coding RNA, most notably miRNA.^8^ Interestingly, miRNA expression in exosomes may be altered based on pathological changes associated with disease states.^9,10^

Using novel techniques, it is possible to enrich for cell-specific EVs from a population of bulk EVs from biofluids. ^7^ In psychiatry and other fields of neuroscience, there has been a keen interest in studying neuron-derived extracellular vesicles that are accessible in plasma. This is done by first extracting total EVs from plasma, and then enriching for neuronal material using an antibody for neural markers such a neural cell adhesion molecule L1CAM.^11^ To further confirm neuronal origin, markers such as miR-9 ^12^as well as cell-specific protein markers, can be used to verify neuronal-enrichment using this method. Recent studies have identified both protein and miRNA cargo of neuron-derived extracellular vesicles to be used as an early biomarker of Alzheimer’s disease and cognitive impairment.^12–14^ Finally, most recently, proteins in L1CAM+ exosomes have been used as a biomarker for MDD ^15^ and bipolar disorder treatment response.^16^

In this study, we investigated neuron-derived extracellular vesicles (NDEVs) and their miRNA cargo as a function of ADT response. We first demonstrated that it is possible to extract NDEVs from plasma and sequence their miRNA cargo. We then characterized NDEVs and showed that their miRNA cargo overlapped with the cargo of EVs obtained directly from brain tissue. Furthermore, our analysis revealed that the predicted targets of the NDEV miRNA cargo were enriched for brain specific mRNA transcripts. Group specific findings indicated that untreated MDD patients had smaller vesicles than controls, and size of NDEV increased with treatment response. Finally, we identified nine NDEV miRNAs with expression profiles that correlated with changes in the Montgomery-Asberg Depression Rating Scale (MADRS) score from baseline to week eight of ADT. Of these, a combination of three miRNAs was highly predictive of ADT response.

## Methods

### Subject details: human plasma

This study consisted of participants who were chosen from a larger antidepressant drug response cohort.^17^ For this study, a total of 60 male and female participants were used: 20 healthy controls and 40 diagnosed with MDD. Samples were matched for age and sex. Patients were ascertained at a community outpatient clinic at the Douglas Mental Health University Institute. All subjects provided informed consent, and the project was approved by The Institutional Review Board of the Douglas Mental Health University Institute. Subjects were excluded if they had comorbidity with other major psychiatric disorders, if they had positive tests for illicit drugs during the study, or general medical illnesses. Control subjects were excluded if they had a history of ADT. Patients had a diagnosis of MDD without psychotic features according to the DSM-IV. After a washout period, MDD patients were given the recommended dosage of escitalopram (Cipralex). After eight weeks of treatment, participants were classified as either escitalopram responders (RES) or non-responders (NRES) using the Montgomery-Asberg Depression Rating Scale (MADRS). Response was determined by a greater than 50% decrease in MADRS score after eight weeks of treatment. Of the patients used in this study, 20 were RES and 20 were NRES. Cohort details can be found in Table 1.Six millilitres of blood were drawn before drug treatment (T0), and eight weeks after drug treatment (T8). The same time points were used for blood draws from untreated controls. Blood was centrifuged at 542g for eight minutes at room temperature to separate plasma from whole blood. Plasma was stored at −80 °C until use. Detailed methods of EV isolation, NDEV enrichment, EV characterization, small RNA isolation, sequencing, and analysis can be found in the supplementary material.

**Table 1.** Antidepressant response cohort demographic information.

|  | Controls<br>(n=20) | MDD-RES<br>(n=20) | MDD-NRES (n=20) |
| --- | --- | --- | --- |
| Number | 20 | 20 | 20 |
| Sex | 6M/14F | 9M/ 11F | 9M/ 11F |
| Age | 45.05 ± 13.76 | 39.30 ± 11.21 | 44.3 ± 10.65<sup>ns</sup> |
| Ethnicity | 20 Caucasian | 20 Caucasian | 19 Caucasian/ 1 no answer |
| Time of blood draw | 9:00 AM | 9:00 AM | 9:00 AM |
| Antidepressant treatment | None | Escitalopram | Escitalopram |
| Fasting | 20Y | 20Y | 20Y |
| MADRS T8 | - | 5.63 ± 4.33 | 26.05 ± 8.05*** |
| MADRS T0 | - | 30.22 ± 5.58 | 33.35 ± 6.19 |
| ΔMADRS (T8/T0) | - | 0.18 ± 0.13 | 0.77 ± 0.19*** |

\*\*\* $p < 0.001$
ns ( $F_2 = 1.29$ , $p = 0.280$ )
MDD-RES= escitalopram responder, MDD-NRES=escitalopram non-responder

### Subject details: human brain

As previously described by Lutz and colleagues^18^, postmortem Anterior Cingulate Cortex (ACC) brain tissue was obtained in collaboration with the Quebec Coroner’s Office and the suicide section of the Douglas-Bell Canada Brain Bank. Ethical approval was obtained from The Institutional Review Board of the Douglas Mental Health University Institute and written informed consent was obtained from the family. Cohort details can be found in Supplementary Table 3. Psychological autopsies were performed by trained clinicians on cases and controls, as previously described.^18^ The control group had no history of suicidal behavior or major mood or psychotic disorders, with diagnoses assigned based on DSM IV. Detailed methods of RNA-sequencing can be found in the supplementary material, and the paper referenced above.

Additionally, one (1) human postmortem brain sample was used from Brodmann area 9 for EV extraction from brain tissue. Sample details can be found in Supplementary Table 4. Detailed methods of EV extraction, small RNA sequencing, and analysis can be found in the supplementary material.

### EV size and quantity analysis

Statistical analysis for EV size and quantity was done using GraphPad Prism 5. Data is displayed as mean ± SEM. D’Agostino & Pearson normality test was used to determine normality for all datasets, and unequal variance was corrected for if present. Outliers were removed, according to ROUT’s test results (Q=1%). Appropriate statistical tests were used for each scenario, as described in figure captions. Generalized linear model correcting for age was used to calculate the regression between both the fold change between time points of EV size (T8/T0) and EV number (T0/T8) against the fold change of MADRS score (T8/T0).

### Sequencing analysis for miRNA from NDEVs

Sequencing reads were trimmed with cutadapt to remove technical sequences. Trimmed reads were submitted to the ERCC’s exceRpt small RNA-Seq pipeline (v.4.6.2) which aligned the reads to the human genome and quantified the various miRNAs into counts. Using DESeq2, the counts were normalized using the “postcounts” methods to account for the large number of zeroes found in the count table. Principal component analysis was carried out to find underlying sources of biases. Each potential covariate was correlated to each of the first five principal components as sorted by decreasing order of explained variance. Only covariates with a significant correlation (p-value<0.05) were included in the statistical model. In the end, a generalized linear model (GLM) controlling for the samples’ sequencing batches, the number of non-zero miRNAs, as well as the sex of each patient was used. Only miRNAs with at least three reads in 70% of one of our six groups of interest (three clinical groups at two different time-points) were kept for further analyses.

### MiRNA expression analysis

A generalized linear model was performed using cases only to identify miRNAs whose fold-change had a significant relationship with changes in MADRS score. The GLM was fitted on the miRNAs’ fold-changes (the expression at T8 divided by the expression atT0) with the outcome variable of the delta MADRS expressed as a ratio (the MADRS at T8 divided by the MADRS at T0). Wald’s test was used to calculate p-values (chi-square based) with a sandwich correction. Those with a p-value of <0.1 were considered for further analysis. A receiver operating characteristic (ROC) curve with backwards stepwise regression analysis was performed to determine if any of those miRNAs with a p<0.1 had predictive value.

## Results

### NDEVs were smaller than EVs from plasma and were reflective of brain tissue

We extracted total plasma EVs and subsequently, NDEVs from MDD patients undergoing a clinical trial of escitalopram, a selective serotonin reuptake inhibitor used for the treatment of depression, as well as untreated psychiatrically healthy controls. Blood samples from 40 MDD patients and 20 controls were randomly selected from a larger cohort to include equal number of samples in each group (N=20, RES, NRES, controls). Cohort details can be found in Table 1. Blood was collected before MDD patients were treated, (T0) and eight weeks after escitalopram treatment (T8). Blood was collected at the same time points for the untreated controls. After eight weeks of drug use, MDD patients with a greater than 50% reduction in MADRS scores were classified as escitalopram responders. (Figure 1A).

**Figure 1.**
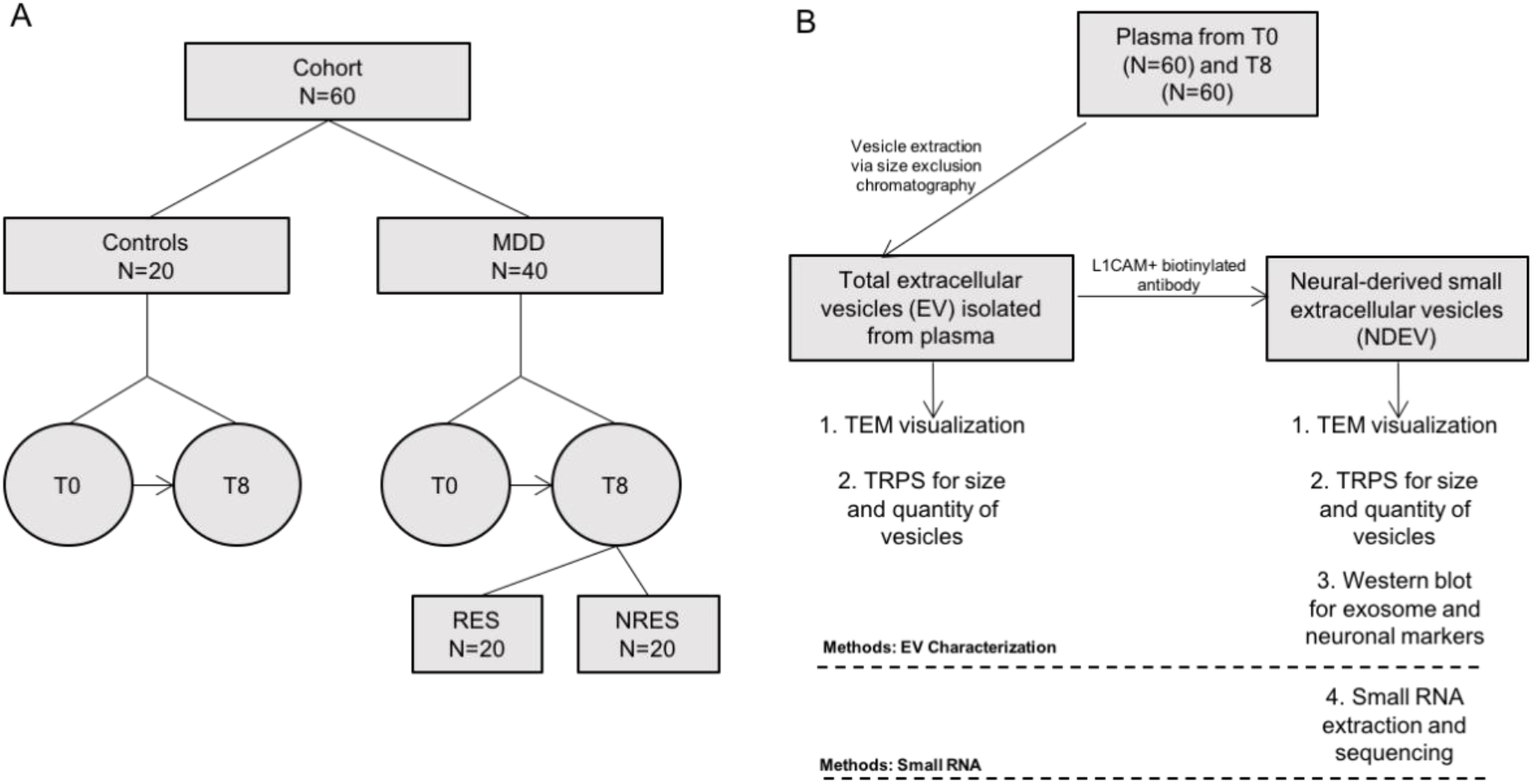
Cohort and Methods. **A.** Our cohort consisted of 20 controls and 40 MDD patients, 20 of which were responders (RES) and 20 non-responders (NRES) following eight weeks of escitalopram treatment. Plasma was drawn before treatment (T0) and eight weeks after (T8) for all groups, with untreated controls being drawn at the same time points. **B.** EVs were extracted from plasma of all groups at T0 and T8. Some of the total plasma EV fraction was kept for characterization, while the rest was used for neuron enrichment using an L1CAM biotinylated antibody. All EVs were characterized using TEM, TRPS, and western blots. RNA extraction and sequencing was done with NDEV fraction only.

We first aimed to characterize NDEVs from plasma, and determine if the vesicles were representative of cells in the brain. To do this, we extracted total EVs from plasma and then further enriched for NDEVs using a biotinylated L1 cell adhesion molecule (L1CAM) antibody (Figure 1B, Supplementary Methods). We performed a rigorous characterization to confirm proper EV extraction and enrichment, including the characterization of both protein and vesicles (Figure 1B). Transmission electron microscopy (TEM) images demonstrated that NDEVs were of expected size and shape (Figure 2A, panel i). Although vesicles are spherical in solution, after the drying process, vesicles have a cup-shaped appearance.^19^ Immunolabeling of NDEVs with CD81 demonstrated that a common exosomal marker was present on the vesicles (Figure 2A, panel ii, white arrows).

**Figure 2.**
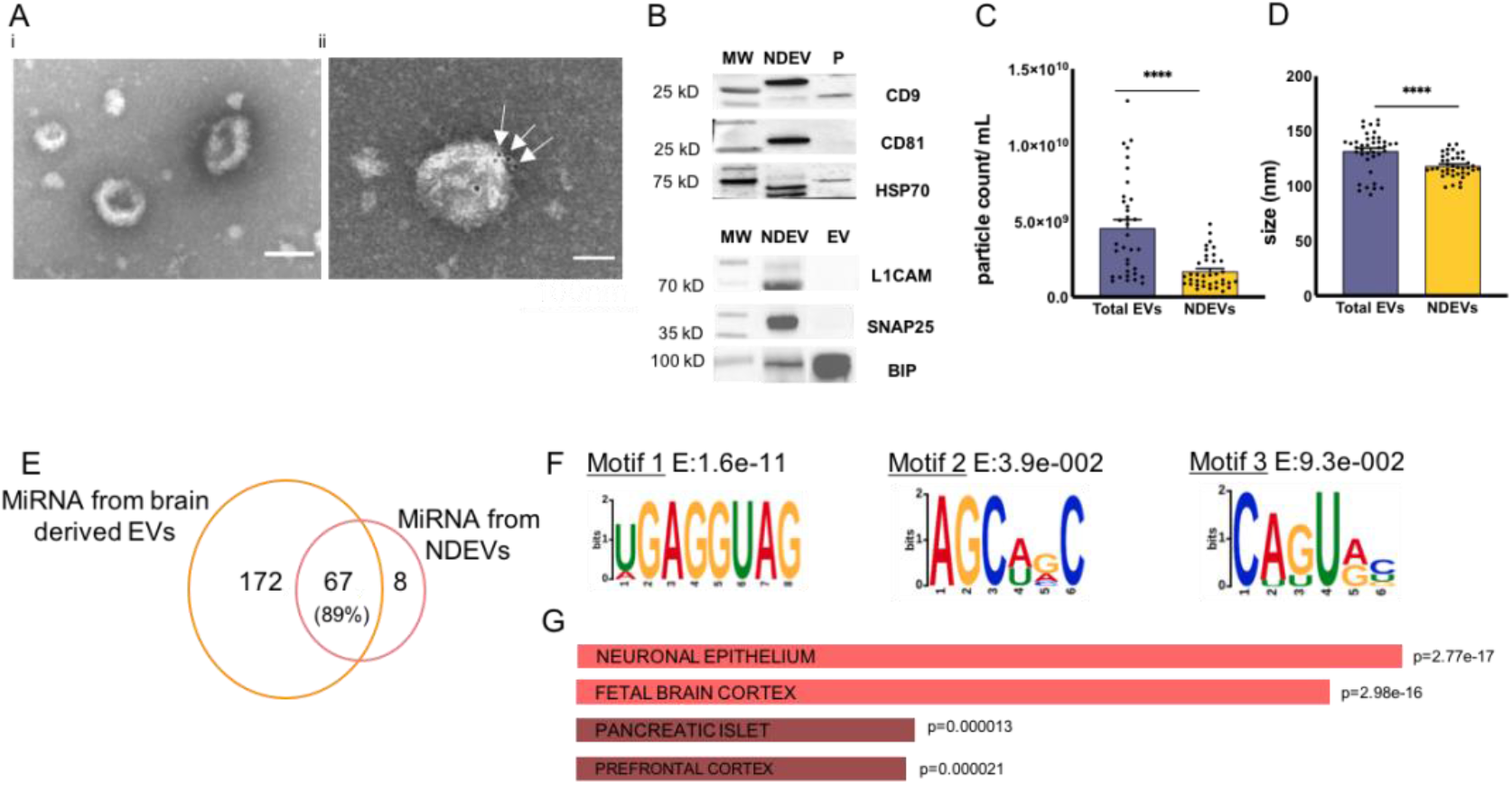
NDEV vesicle and cargo characterization. **A.** TEM images of NDEVs, second panel including immunolabelling of CD81. Scale bars =100nm **B.** Protein markers for EVs and neuron markers within our EV fractions. MW= molecular weight markers NDEV= neuron-derived EVs; EV= total plasma EVs extracted from plasma (pre-neuron enrichment) P= protein fraction collected after EV fraction during size exclusion chromatography (void of extracellular vesicles). The bottom western blot was rearranged for ease of viewing; Supplementary Figure 1 contains the original blot. **C.** On average 4.53e9 (± 3.23e9) particles/mL were extracted from plasma. Our NDEV fraction contained approximately 1.68e9 (± 1.13e9) particles/mL. (N_Total_=35, N_NDEV_= 35, Wilcoxon matched pairs signed rank test, two tailed, p<0.0001, W=-519.0) **D.** Sizing results showed that the average size of NDEVs (118.8 ± 10.07nm) was smaller than total plasma EVs extracted from plasma (131.8 ± 18.46nm). (N_Total_=40, N_NDEV_=40, paired t-test, two tailed, df=39, t=4.583, p<0.0001). **E.** 89% of our list of miRNAs within NDEV overlapped with miRNAs present in brain tissue. **F.** Three consensus binding motifs were significantly enriched within our list of miRNAs from NDEV. **G**. Tissue expression of the genes that were predicted targets of all three motifs showed enrichment for genes in brain tissue. Coral= p<10e-10, brown = p<0.05. All data in this figure is represented as mean ± SEM. ****p<0.0001

We tested common exosomal markers, CD63 and CD9, and heat shock protein 70 (HSP70), another protein frequently found in exosomes, on our EV samples. All were present in the NDEV fraction, and were either not present or depleted in a fraction void of exosomes (Figure 2B). We observed a strong presence of synaptosome associated protein 25 (SNAP25), a marker of brain neurons, in our NDEV fraction, which was not detectable in our total EV fraction (Figure 2B, Supplementary Fig.1). As expected,^20^ we observed cleaved L1CAM (between 70-100kD) enriched in the NDEV fraction (Figure 2B, Supplementary Fig.1). Finally, binding immunoglobulin protein (BiP), a marker for endoplasmic reticulum vesicle contamination, was minimal after NDEV enrichment (Figure 2B, Supplementary Fig.1).

Tunable resistive pulse sensing (TRPS) was used to assess the size and quantity of EVs in our samples. A subset of our samples from each group was randomly chosen for TRPS; and both total plasma EVs and NDEVs were analyzed at each time point. On average 4.53e9 (± 3.23e9) particles/mL were extracted from plasma, and our NDEV fraction contained approximately 1.68e9 (± 1.13e9) particles/mL (Figure 2C). On average, NDEVs (118.8 ± 10.07nm) were smaller than total EVs extracted from plasma (131.8 ± 18.46nm, p<0.0001) (Figure 2D).

As exosome cargo is primarily composed of miRNA^8^, we built small RNA libraries for sequencing to characterize the miRNA cargo of NDEV. Given the low abundance of individual miRNAs in EV cargo, we focused on miRNAs with at least three counts in 70% of a given group (controls, RES, and NRES) as we reasoned they would be markers of the most active and biologically relevant processes. After applying this threshold, 75 miRNAs were considered for further analyses (Supplementary Table 1).

Since we expected NDEV cargo to mirror the cargo of EVs extracted from brain tissue, we also extracted EVs from a human cortical sample (Brodmann area (BA) nine). We observed that 67 of the 75 miRNAs (89%) within NDEVs were also present in EVs from brain tissue (Figure 2E). This is in contrast with the poor concordance observed when comparing NDEV cargo with that of EVs from other origins. For instance, we did not observe any overlap of miRNA species between our NDEV cargo list and a list of miRNAs extracted from exosomes in human saliva (Supplementary Figure 2).^21^

To further investigate the relevance of the NDEV miRNAs in the brain, we performed a consensus motif enrichment discovery analysis using MEME.^22^ Focusing on the seed regions of the miRNAs, we detected three consensus binding motifs enriched within miRNAs from NDEV (E<0.1) (Figure 3B.). Using ENSEMBL,^23^ we generated a list of 1,760 genes that had predicted binding sites for all three of these motifs within their 3’UTR. Using the enrichment analysis tool Enrichr ^24^, and tissue expression atlas ARCHS4^25^, the predicted gene targets for all three motifs were shown to be enriched in neuronal epithelium (p=2.77e-17), fetal brain cortex (p=2.98e-16), pancreatic islet (p=0.000013), and prefrontal cortex tissue (p=0.000021) (Figure 2G).

**Figure 3.**
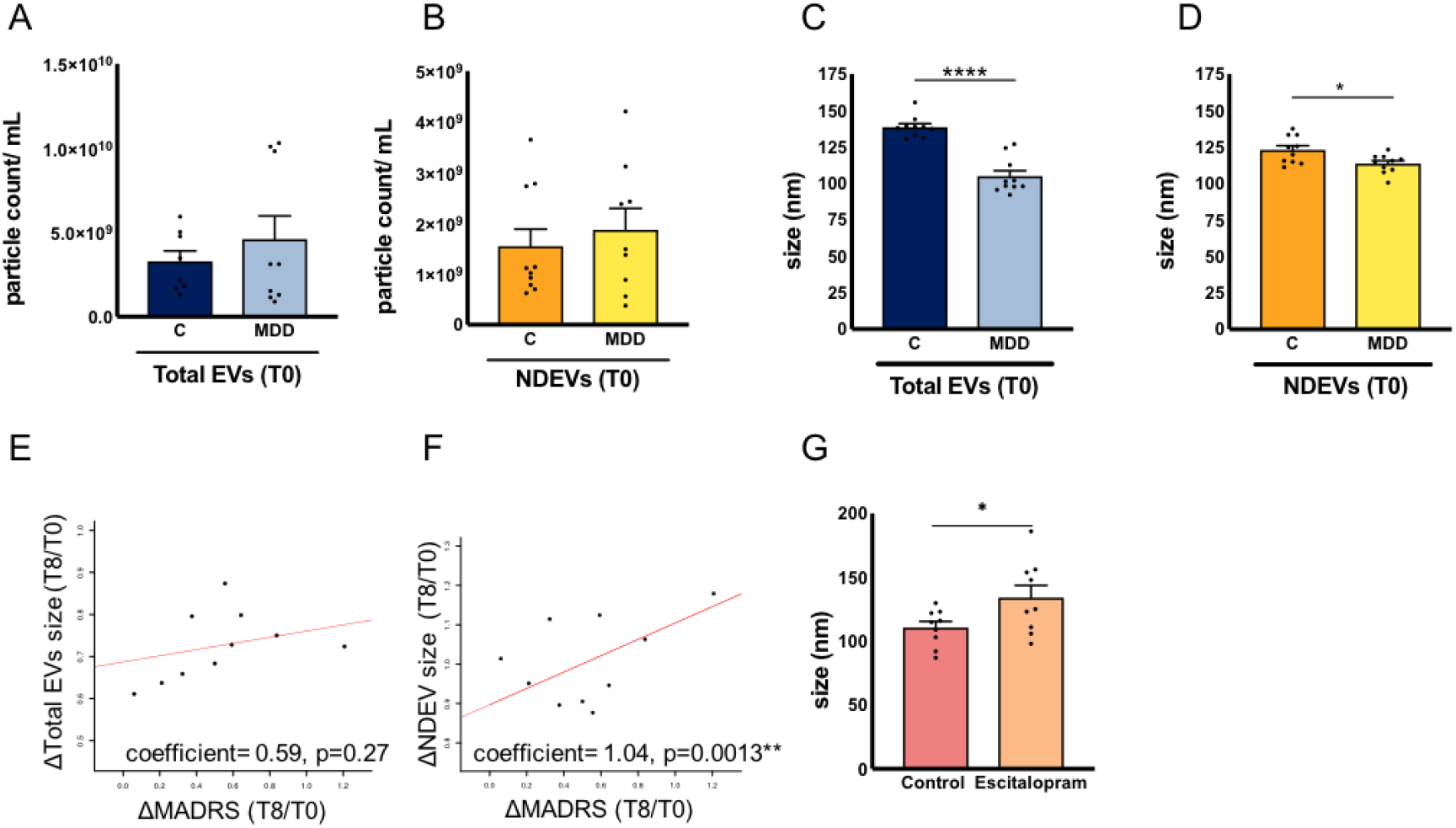
NDEV size was altered following antidepressant drug response. **(A)** At T0, there were no differences between the number of total plasma EVs between MDD patients (4.59e9 ± 4.18e9) and controls (3.28e9 ± 1.77e9) (N_C_=8, N_MDD_=9 unpaired two tailed t-test with Welch’s correction for unequal variances, df=11.05, t=0.8577, p=0.4093). **(B)** At T0 there were no differences between number of NDEV between MDD patients (1.87e9 ± 1.27e9) and controls (1.54e9 ± 1.07e9) (N_C_=10, N_MDD_=9 Mann-Whitney test, U=40, p=0.7197). However, at T0, **(C)** total plasma EVs of MDD patients (104.9 ± 12.12nm) were smaller than those from psychiatrically healthy controls (138.7 ± 7.24nm) (N_C_=10, N_MDD_=10, Mann-Whitney test, two-tailed U=0, p<0.0001). Furthermore, NDEV **(D)** were smaller in MDD patients (113.6 ± 6.5nm) compared to controls (123.0 ± 9.40nm) (N_C_=10, N_MDD_=10, unpaired, two tailed, t-test with equal variance, df=18, t=2.599, p=0.0181). **E.** Regression between changes in MADRS score between T8 and T0, and the size of total plasma EVs showed no significant relationship (coefficient= 0.59, p=0.27).**F.** Regression between changes in MADRS score between T8 and T0, and changes in NDEV size showed a positive relationship, indicating an increase in NDEV size with response (coefficient= 1.04, p=0.0013) **G.** EVs released from treated HEK293 cells (134.1 ± 28.71nm) had larger EVs compared to untreated controls (110.8 ± 14.45nm) (N_C_=9, N_escitalopram_=9, unpaired two tailed t-test, df=16, t=2.178, p<0.05). All data is represented as mean ± SEM. *p<0.05,****p<0.0001

### Neuron-derived EVs demonstrated changes during antidepressant treatment as a function of drug response

Given the relevance of NDEV cargo to brain tissue, and the differences in sizes between NDEV and total plasma EVs, we wanted to examine the dynamic changes of NDEVs that occur during ADT. By comparing variations in NDEVs both at T0 and across time in escitalopram RES, NRES, and untreated controls, we assessed alterations that were specific to a given group.

Before drug treatment (T0), we found no differences in the number of total plasma EVs (Figure 3A) or NDEV (Figure 3B) between MDD patients and controls. To identify whether changes in EV number during drug treatment were related to changes in MADRS score (response), a logistic regression correcting for age was performed. There was no significant relationship between a change in MADRS score and change in the number of total plasma EVs (Supplementary Figure 3A), or NDEVs (Supplementary Figure 3B).

Analysis of EV size before treatment found that both total plasma EVs (Figure 3C, p<0.0001) and NDEVs (Figure 3D, p<0.05) of MDD patients were smaller than controls. Regression analysis found no significant relationship between a change in total plasma EV size and a change in MADRS score during eight weeks of treatment (Figure 3E). However, there was a strong positive relationship between a change in MADRS score and a change in NDEV size with treatment, suggesting that NDEV size increases with treatment response (coefficient= 1.04, p<0.01) (Figure 3F). To test the effect of escitalopram on EV size, we treated HEK293 cells with escitalopram. The average size of EVs released from treated cells was larger than those of untreated cells (p<0.05) (Figure 3G).

Next, we aimed to investigate how NDEV cargo was affected during ADT. We performed a logistic regression on MDD cases to investigate if any of the 75 NDEV miRNA cargo previously identified were altered with treatment response. The analysis revealed that nine miRNAs changed over time (p<0.1) after sandwich correction: miR-423-3p, miR-191-5p, miR-486-5p, miR-30d-5p, miR-425-5p, miR-25-3p, miR-21-5p, miR-335-5p, and miR-126-5p (Table 2). We then used a backwards stepwise regression analysis to determine if any combination of the nine miRNAs were predictive of response. The combination of miR-21-5p, miR-30d-5p, and miR-486-5p changes over antidepressant treatment associated with ADT response (AUC:0.8254, specificity: 84%, sensitivity: 76%, p<0.01) (Figure 4A).

**Table 2.** MiRNA altered across time as a function of changes in MADRS score. Logistic regression analysis of MDD cases demonstrated that nine miRNAs were altered across time as a function of change in MADRS score (p-value with sandwich correction <0.1). ROC curve analysis with these nine miRNAs demonstrated that miR-21-5p, miR-30d-5p, and miR-486-5p are predictive of antidepressant drug response (specificity 84%, sensitivity 76%, F=4.355, p<0.01), as outlined in Figure 4A.

| miRNA | coefficient of<br>expression with<br>ΔMADRS | p value |
| --- | --- | --- |
| hsa-miR-423-3p | -0.0646 | 0.004 |
| hsa-miR-191-5p | -0.032 | 0.01 |
| hsa-miR-486-5p | -0.0389 | 0.021 |
| hsa-miR-30d-5p | -0.0472 | 0.035 |
| hsa-miR-425-5p | -0.0298 | 0.043 |
| hsa-miR-25-3p | -0.0305 | 0.051 |
| hsa-miR-21-5p | 0.0232 | 0.067 |
| hsa-miR-335-5p | -0.0299 | 0.072 |
| hsa-miR-126-5p | 0.0351 | 0.074 |

**Figure 4.**
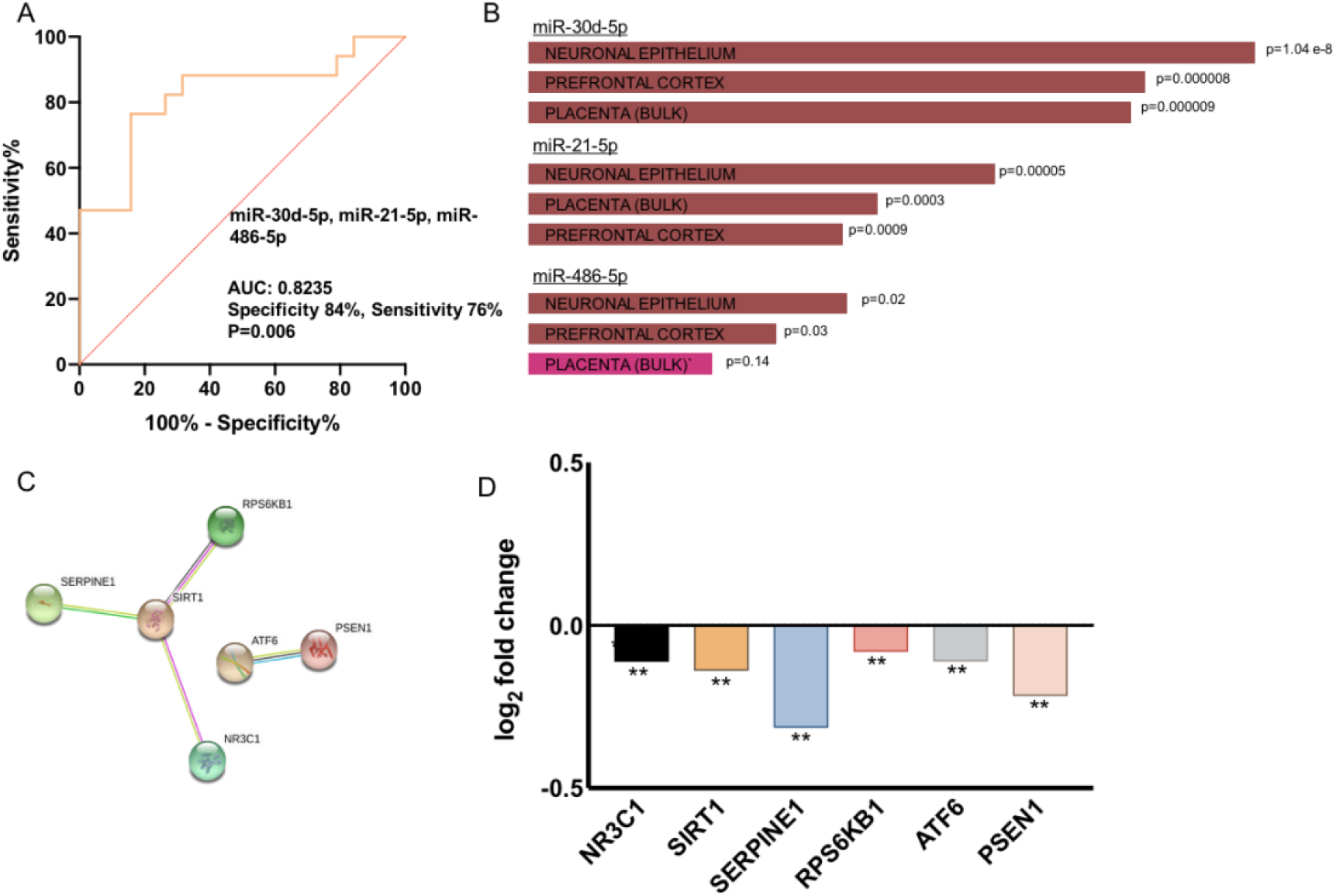
NDEV miRNA cargo was altered in response to ADT, with some cargo predicting response. **A.** Backwards step regression identified miR-21-5p, miR-30d-5p, and miR-486-5p to be predictive of ADT response (specificity 84%, sensitivity 76%, F=4.355, p<0.01) **B.** MiR-21-5p, miR-30d-5p, and miR-486-5p were enriched in brain tissue. The three most enriched tissues per Enrichr/ARCHS4 are shown: brown (p<0.05), fuchsia (n.s) **C.** Protein-protein interactions of the 25 predicted targets of miR-486-5p that were the most significantly altered in the ACC of depressed individuals who died by suicide. Genes excluded were not part of any interaction. **D.** Fold change of altered genes in the brain that were part of the network in Figure 4C.(N_C_=26, N_MDDS_=27, differential expression was analyzed using DESeq2, **p <0.01

A target prediction analysis of miR-21-5p, miR-30d-5p, and miR-486-5p generated lists of 441, 594, and 391 targets, respectively. Targets for all miRNAs were enriched in brain tissue (Figure 4B), and for pathways including MAPK signaling and axon guidance (Supplementary Figure 4). As the predicted targets of NDEV miRNAs are enriched in the brain, we wondered whether these targets showed comparable changes within the anterior cingulate cortex (ACC) of individuals who died by suicide during an episode of major depression. Given that these individuals did not respond to ADT, we expected changes to be opposite from those of ADT responders. We focused on the predicted gene targets from each of the miRNAs that overlapped with genes that were significantly altered between controls and those who died by suicide in the brain cohort. This resulted in a total of 174 targets: 113 (miR-30d-5p), 14 (miR-21-5p), and 47 (miR-486-5p) (p<0.05).

STRING analysis was performed on the top 25 predicted targets that overlap with changes found in the brain (except for miR-21-5p where there was only a list of 14 targets). The targets of miR-486-5p showed significant functional interactions for "cellular response to drug" (FDR=0·007), regulation of signal transduction (FDR=0·02), and "activation of MAPKK activity" (FDR= 0·04) (Supplementary Table 2). Genes that interacted within this network are displayed in Figure 4C, and the expression of these genes in the depressed ACC is displayed in Figure 4D.

## Discussion

Our examination of EVs from plasma of individuals undergoing ADT revealed a novel, minimally-invasive way of assessing molecular changes associated with the efficacy of ADT. Given that our findings were from EVs with a neuron-specific surface marker, we may extrapolate our results to changes occurring directly in the brain. This technique provides a promising tool that gives us insight into potential mechanisms and pathways that are altered in the brain.

Our findings suggest that EVs from untreated MDD patients are smaller than those of controls, and that this size difference can be reversed by ADT. Although our size results are interesting, and the effects of escitalopram on neurons expressing the serotonin transporter are a possible and enticing hypothesis to explain NDEV size changes, there is limited information on mechanisms that could explain the size differences observed. It is well documented that exosomes released from even the same cell type are heterogeneous.^26^ An alternative hypothesis is that antidepressants affect EV release and biogenesis, and could potentially stimulate a greater release from a given cell type, or alter the release of exosomes altogether. Current evidence points to two main pathways of exosome biogenesis, one dependent on endosomal sorting complex required for transport (ESCRT) proteins, the other, ESCRT-independent.^27^ Limited evidence suggested that when vital proteins involved in the ESCRT-dependent pathway are blocked, larger exosomes are secreted.^26^ Our last hypothesis is that smaller exosomes simply contain less cargo. This could suggest an overall reduction of communication via exosome cargo in untreated MDD patients, which may be rescued upon treatment response to escitalopram.

Our study also suggested that changes in expression of a combination of three miRNAs within NDEV were predictive of ADT response. This is intriguing, as miRNAs often work together to downregulate targets and pathways. ^28^ The three NDEV miRNAs identified in this study are predicted to regulate the MAPK pathway, which has been previously implicated in ADT response, including in a study that isolated miRNAs from plasma samples.^4^ Individually, all three miRNAs identified have been associated with MDD and/or ADT response. MiR-21-5p was identified as a biomarker that discriminated between duloxetine remitters and non-remitters.^29^ Interestingly, miR-21-5p was shown to target cell adhesion molecule L1 like (CHL1), which is an important regulator of brain development and maintenance of neural circuits.^30^ MiR-30d-5p was associated with late-life depressive symptoms, and is an important regulator in the brain. ^31^ Finally, miR-486-5p was downregulated in the brains of those who died by suicide and was a “hub” connecting to many other miRNAs that were also downregulated in the brain.^32^ Of note, one target of miR-486-5p that was present in our network analysis is nuclear receptor subfamily 3 group C member 1 (NR3C1), the gene that codes for the glucocorticoid receptor. NR3C1 has been extensively implicated in MDD through hypothalamic-pituitary-adrenal axis dysregulation.^33^

Although our study provides a promising avenue for investigating molecular changes associated with psychiatric illnesses, it is not void of limitations. An important limitation is that the antibody used to isolate NDEV cannot separate peripheral from central neurons. Although our NDEVs were enriched for SNAP25 and had high miRNA content overlap with EVs extracted from brain tissue, L1CAM is a general neuronal marker. Future studies should consider using an antibody specific for central neurons only. Further studies should also be completed investigating EVs from brain tissue. Given that miRNA are already quite stable to post-mortem degradation ^34,35^, and that EVs may provide extra protection for these miRNAs in the brain, EVs may be intriguing candidate biomarkers that may remain useful after death. Another limitation of this study is the relatively small sample size. As technology advances and investigations of NDEVs become less time consuming, larger sample sizes should be investigated.

In summary, this study provides direct insight into molecular changes in the brain through the use of plasma extracted NDEVs. This population of EVs has the potential to act as a clinically accessible marker of MDD and ADT response. Given the inherent challenges in sampling CNS tissue from living subjects, this work provides an encouraging avenue to study the molecular mechanisms contributing to neuropsychiatric illnesses and treatment response. Nonetheless, further research is required to replicate and expand these findings.

## Acknowledgements

Funding for this study was provided by grants to GT from the Canadian Institute of Health Research (CIHR) (FDN148374 and EGM141899), and by the *Fonds de recherche du Québec* - *Santé* (FRQS) through the Quebec Network on Suicide, Mood Disorders and Related Disorders. In addition, GT holds a Canada Research Chair (Tier 1) and a NARSAD Distinguished Investigator Award. CAN-BIND is an Integrated Discovery Program carried out in partnership with, and financial support from, the Ontario Brain Institute, an independent non-profit corporation, funded partially by the Ontario government. The opinions, results and conclusions are those of the authors and no endorsement by the Ontario Brain Institute is intended or should be inferred. Funding support was also provided by a grant from the Ontario Research Fund: Research Excellence (RE-08-027) SD. The Douglas-Bell Canada Brain Bank is funded by a Healthy Brains for Healthy Lives (CFREF) Platform Grant to GT and NM and by the *Réseau Québécois sur le suicide, le troubles de l’humeur et les troubles associés* (FRQS). SS is funded by the Canadian Institute of Health Research (CIHR). We would like to thank Dr. David Junker and his lab, especially Philippe DeCorwin-Martin, for the use and assistance of the qNano. We would also like to thank the entire Turecki lab for their feedback and guidance.

## Author Contributions

Conceptualization, S.S., C.N., and G.T. Methodology, S.S., C.N., M.W., and J.F.T. Formal Analysis, S.S. and J.F.T. Investigation, S.S., C.N., M.W., P.I., L.M.F. and J.Y. Writing-original draft S.S., C.N., and G.T. Writing-review and editing J.F.T., L.M.F., P.I., S.R., S.K., J.A.F., and N.M. Resources, S.R., S.K., J.A.F., N.M., and G.T. Funding Acquisition, G.T, S.R., S.K., J.A.F., N.M.

